# Rethinking Remdesivir: Synthesis, Antiviral Activity and Pharmacokinetics of Oral Lipid Prodrugs

**DOI:** 10.1101/2020.08.26.269159

**Authors:** Robert T. Schooley, Aaron F. Carlin, James R. Beadle, Nadejda Valiaeva, Xing-Quan Zhang, Alex E. Clark, Rachel E. McMillan, Sandra L. Leibel, Rachael N. McVicar, Jialei Xie, Aaron F. Garretson, Victoria I. Smith, Joyce Murphy, Karl Y. Hostetler

**Author notes:** Corresponding Authors: Robert T. Schooley, Karl Y. Hostetler. These authors contributed equally to the manuscript (Robert T. Schooley, Aaron F. Carlin).

## Abstract

Remdesivir (RDV, GS-5734) is currently the only FDA-approved antiviral drug for the treatment of SARS CoV-2 infection. The drug is approved for use in adults or children 12-years or older who are hospitalized for the treatment of COVID-19 on the basis of an acceleration of clinical recovery for inpatients with this disease. Unfortunately, the drug must be administered intravenously, restricting its use to those requiring hospitalization for relatively advanced disease. RDV is also unstable in plasma and has a complex activation pathway which may contribute to its highly variable antiviral efficacy in SARS-CoV-2 infected cells. Potent orally bioavailable antiviral drugs for early treatment of SARS-CoV-2 infection are urgently needed and several including molnupiravir and PF-07321332 are currently in clinical development. We focused on making simple, orally bioavailable lipid analogs of Remdesivir nucleoside (RVn, GS-441524) that are processed to RVn-monophosphate, the precursor of the active RVn-triphosphate, by a single-step intracellular cleavage. In addition to high oral bioavailability, stability in plasma and simpler metabolic activation, new oral lipid prodrugs of RVn had submicromolar anti-SARS-CoV-2 activity in a variety of cell types including Vero E6, Calu-3, Caco-2, human pluripotent stem cell (PSC)-derived lung cells and Huh7.5 cells. In Syrian hamsters oral treatment with ODBG-P-RVn was well tolerated and achieved therapeutic levels in plasma above the EC_90_ for SARS-CoV-2. The results suggest further evaluation as an early oral treatment for SARS-CoV-2 infection to minimize severe disease and reduce hospitalizations.

## INTRODUCTION

Over the past 18 years, spillover events have introduced the highly transmissible beta-coronavirus strains SARS CoV, MERS CoV, SARS CoV-2 into the human population.[1–3] Although case fatality ratios have varied, each virus induces substantial morbidity and mortality – especially among those over 55 and/or those with underlying co-morbid medical conditions.[4,5] Although SARS CoV and MERS CoV were largely contained by epidemiological interventions, the most recent 2019 outbreak has evolved into a global pandemic responsible for over 160 million infections and over 3.5 million deaths.[6] With over 30 million cases and nearly 600,000 deaths at this writing, the US remains in the center of the pandemic. Intensive and economically disruptive social distancing measures have blunted several viral surges, but this approach is not sustainable and infection rates have increased each time restrictions have eased. [7] Fortunately, several highly effective vaccines have emerged over the past six months and their distribution in the US and many other high-income countries has had a major impact on SARS CoV-2 associated morbidity and mortality. [8,9] Despite these highly noteworthy successes in vaccine development, gaps and delays in global vaccine delivery, the emergence of viral variants against which vaccine protection is compromised and a relatively sizable immunocompromised population who are unable to fully respond to vaccination make it clear that infections will continue and that highly effective antiviral therapy is required.

Remdesivir nucleoside triphosphate (RVn triphosphate) potently inhibits enzymatic activity of the polymerase of every coronavirus tested thus far, including SARS CoV-2. [10–13] The drug has recently been approved by the FDA for the treatment of adults and children aged 12 or over who are hospitalized for COVID-19. [14] This broad activity reflects the relative molecular conservation of the coronavirus RNA-dependent RNA polymerase (RdRp). Remdesivir (RDV) is an aryloxy phosphoramidate triester prodrug that must be converted by a series of reactions to RVn triphosphate, the active antiviral metabolite. (Fig 1) Although RVn-triphosphate is an excellent inhibitor of the viral RdRp [11,15], RDV’s antiviral activity is highly variable in different cell types which may be due to variable expression of the four enzymes required for conversion to RVn-P [16]. RDV’s base is a 1’-cyano-substituted adenine C nucleoside (GS-441524, RVn) that is thought to be poorly phosphorylated. To bypass the perceived slow first phosphorylation, the developers relied on an aryloxy phosphoramidate triester prodrug that is converted by a complex series of four reactions to remdesivir nucleoside monophosphate (RVn-P) that is then efficiently converted to RVn triphosphate, the active metabolite. RDV may be more active in some SARS-CoV-2 infected tissues than in others, a possible reason for its incomplete clinical impact on SARS-CoV-2. A recent report suggests that some tissues express low levels of the four enzymes that activate RDV in some tissues may be responsible for tissue-specific differences in antiviral activity. [13] Yan and Muller have recently published a detailed analysis of the potential weaknesses of Remdesivir and suggested that RVn (GS-441524) might be a preferable therapy [16]. Remdesivir has beneficial antiviral and clinical effects in animal models of coronavirus infection. [17,18] These effects are primarily demonstrable when administered before or very soon after viral challenge. However, RDV is not highly bioavailable following oral administration and must be administered intravenously, functionally limiting its clinical application to hospitalized patients with relatively advanced disease. It would be clinically useful to have a highly active, orally bioavailable analog of RVn which provides sustained levels of intact antiviral drug in plasma since RDV persistence in plasma is known to be brief. In monkeys treated with intravenous RDV, the plasma level declined by roughly two logs 2 hrs after the infusion ended. [16, 19] In two patients with Covid-19 treated with intravenous RDV, 1 hour after the intravenous infusion stopped a drop of >90% was observed [20]

**Figure 1.**
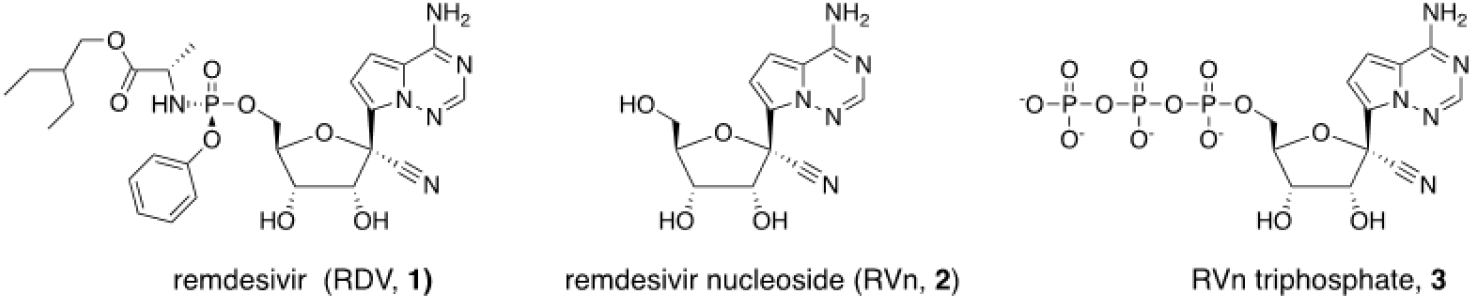
Structures of Remdesivir and related intermediates

Here we report the synthesis and antiviral evaluation of three novel lipid prodrugs of RVn-monophosphate that are active at submicromolar concentrations against SARS-CoV-2 infection in a variety of cell types including Vero E6, Calu-3, Caco-2, PSC-derived human lung cells and Huh7.5 cells. These compounds are stable in human plasma in contrast to Remdesivir and are orally bioavailable as predicted by our prior work with other antivirals of this general design. [21,22] These oral Remdesivir nucleoside phosphate prodrugs could allow earlier and more effective treatment at the time of diagnosis of SARS-CoV-2 infection. In addition, one of these prodrugs represents an approach that may be capable of delivering the antiviral to the lung and away from the liver, the site of Remdesivir’s dose-limiting toxicity, due to its route through intestinal lymph bypassing the portal vein and the liver. [23,24]

## RESULTS

### Synthesis of RVn monophosphate prodrugs

We synthesized the hexadecyloxypropyl-, octadecyloxyethyl- and 1-O-octadecyl-2-O-benzyl-sn-glyceryl-esters of RVn-5’-monophosphate. Compounds **5a**-**5c** were synthesized as shown in Figure 2. Analyses by NMR, ESI mass spec and HPLC were consistent with each structure and demonstrated purities of > 95%.

**Figure 2.**
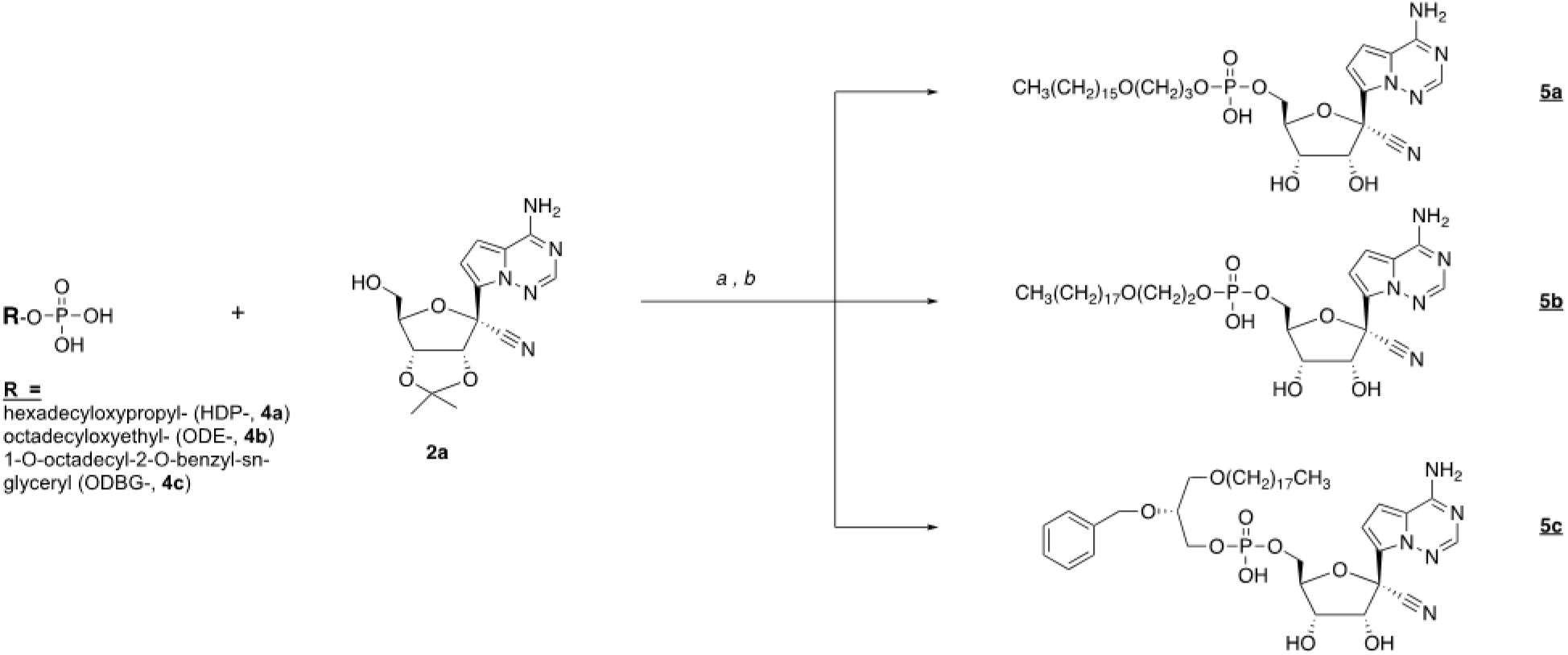
Synthesis of antiviral prodrugs **5a** – **5c**. *Reagents:* a) 2’,3’-isopropylidene RVn (**2a**), DCC, DMAP, pyridine, 90 °C, 24-72 h; b) 37% HCl, THF, 3-18h.

### ODBG-P-RVn stability in human plasma

One of the disadvantages of Remdesivir is instability in plasma where it has been reported to persist for less than 2 hours after intravenous infusion in Rhesus monkeys [13,16,19] and in Covid-19 patients. [20]. We tested the stability of ODBG-P-RVn in human plasma with either K_2_EDTA or sodium heparin (NaHep) as anticoagulant. ODBG-P-RVn was added to human plasma at 37° C for 24 hours and ODBG-P-RVn levels measured at indicated times by LC/MS/MS (Figure 3). This demonstrates that ODBG-P-RVn is stable in human plasma at 37° C for at least 24 hours.

**Figure 3.**
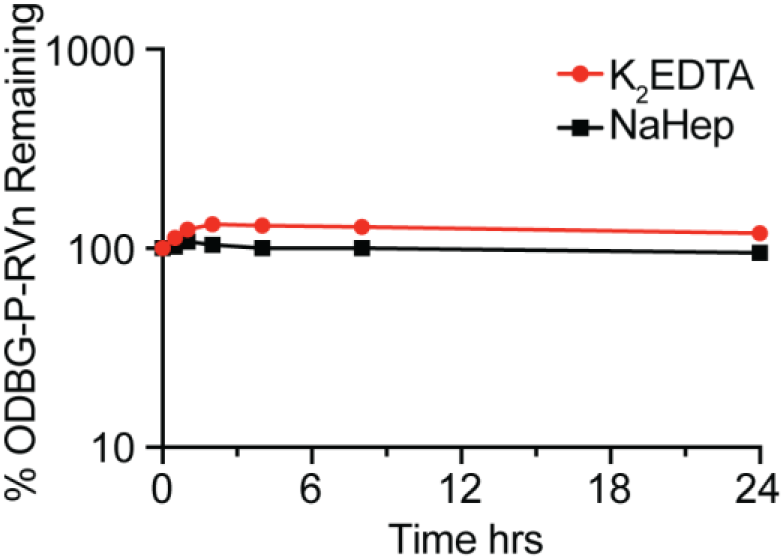
ODBG-P-RVn stability in Human Plasma at 37° C. Plasma was spiked with 2 micrograms/ml concentrations of ODBG-P-RVn and incubated at 37° C for 24 hours. Samples were taken at 0.5, 1,2, 4, 8 and 24 hours and frozen for later analysis by LC/MS/MS.

### Lipid RVn monophosphate prodrugs are potent inhibitors of SARS-CoV-2

To determine if lipid RVn monophosphate prodrugs inhibit SARS-CoV-2 RNA replication, we performed antiviral assays using a clinical isolate of SARS-CoV-2 (2019-nCoV/USA-WA1/2020) in multiple cell lines: African green monkey kidney cells (Vero E6), human PSC-derived lung (PSC-lung) cells, human lung epithelial cell line Calu-3, human colonic epithelial cell line Caco-2, and human hepatocyte cell line Huh7.5. Each cell line was treated with the indicated compound for 30 minutes prior to infection and incubated for 48 hours post-infection. Intracellular viral RNA was measured by quantitative reverse transcription polymerase chain reaction (qRT-PCR). In all cell lines, there was a dose-dependent inhibition of viral RNA by ODBG-P-RVn, ODE-P-RVn, HDP-P-RVn, Remdesivir (RDV) and Remdesivir nucleoside (RVn) (Figure 4A). In Vero E6 cells, the average half-maximal effective concentration (EC_50_) and average 90% effective concentration (EC_90_) of ODBG-P-RVn was 0.14μM and 0.16μM, respectively (Figure 4B and Table 1). The EC_50_ of ODBG-P-RVn in Vero E6 cells was significantly lower than RDV (Table 1). ODE-P-RVn and HDP-P-RVn were also potently antiviral with EC_50_ values of 0.3μM and 0.63μM in Vero E6 (Figure 4B and Table 1). The EC_50_ of ODBG-P-RVn and ODE-P-RVn were less than 0.35μM in PSC-lung and Calu-3, both models of human lung infection (Figure 4B, Table 1). The antiviral activites of ODBG-P-RVn and ODE-P-RVn were significantly better than RVn in PSC-lung cells (Table 1). ODBG-P-RVn, ODE-P-RVn and HDP-P-RVn demonstrated strong antiviral activity in Huh7.5 cells with EC_50_ less than 0.2μM that was not significantly different from RDV or RVn (Figure 4B and Table1). In the Caco-2 cell line, the EC_50_ of ODBG-P-RVn was 0.3μM which was significantly lower than RVn but similar to RDV (Figure 4B and Table 1). In the same cell line, the EC_50_ of ODE-P-RVn was 0.77μM, which was significantly higher than RDV (Figure 4B and Table 1).

**Table 1.** Antiviral Activity, Cytotoxicity and Selectivity of the Compounds.

| Compound | EC<sub>50</sub> (μM) | EC<sub>90</sub> (μM) | CC<sub>50</sub> (μM) | Selectivity | p value EC<sub>50</sub> vs RDV, RVn |
| --- | --- | --- | --- | --- | --- |
| **Vero E6 cells** |  |  |  |  |  |
| RDV | 1.13 | 7.05 | 101 | 89.4 | - |
| RVn | 0.38 | 0.77 | >100 | >263 | - |
| HDP-P-RVn, | 0.63 | 0.73 | >100 | >158 | NS |
| ODE-P-RVn, | 0.30 | 0.33 | >100 | >333 | NS |
| ODBG-P-RVn | 0.14 | 0.16 | 97.9 | 699 | 0.010, 0.311 |
| **PSC-human lung cells** |  |  |  |  |  |
| RDV | 0.14 | 0.23 | 32.7 * | 234 | - |
| RVn | 0.74 | 2.62 | >100 * | >135 | - |
| HDP-P-RVn * | 0.35 | 0.94 | ND |  | - |
| ODE-P-RVn | 0.22 | 0.70 | >100 * | >454 | 0.791, 0.006 |
| ODBG-P-RVn | 0.15 | 0.26 | 61.5 * | 410 | >0.999, 0.002 |
| **Calu-3 cells** |  |  |  |  |  |
| RDV | 0.23 | 0.31 | >100 | >434 | - |
| RVn | 0.15 | 0.18 | >100 | >666 | - |
| ODE-P-RVn | 0.34 | 0.64 | 98.7 | 290 | NS |
| ODBG-P-RVn | 0.30 | 0.33 | 98.2 | 327 | NS |
| **Huh7.5 cells** |  |  |  |  |  |
| RDV | 0.06 | 0.12 | 15.2 | 253 | - |
| RVn | 0.32 | 0.73 | >100 | >312 | - |
| HDP-P-RVn | 0.19 | 0.40 | >100 | >526 | NS |
| ODE-P-RVn | 0.19 | 0.37 | >100 | >526 | NS |
| ODBG-P-RVn | 0.14 | 0.15 | 62.9 | 449 | NS |
| **Caco-2 cells** |  |  |  |  |  |
| RDV | 0.17 | 0.28 | >100 | >588 | - |
| RVn | 0.96 | 1.75 | >100 | >104 | - |
| ODE-P-RVn | 0.77 | 1.25 | >100 | >129 | 0.007, 0.971 |
| ODBG-P-RVn | 0.30 | 0.33 | 88.4 | 295 | 0.968, 0.007 |

**Abbreviations:** RDV, Remdesivir (GS-5734); RVn, Remdesivir nucleoside (GS-441524); HDP-P-, hexadecyloxypropyl-P-; ODE-P-, octadecyloxyethyl-P-; ODBG-P-, 1-O-octadecyl-2-O-benzyl-glycero-3-P-; EC<sub>50</sub>: half-maximal effective concentration; CC<sub>50</sub>: 50% cytotoxic concentration, Selectivity index, CC<sub>50</sub>/EC<sub>50</sub>; statistical analysis comparing LogEC<sub>50</sub> values from separate experiments by one-way ANOVA. CC<sub>50</sub> results by CellTiter-Glo. All experiments performed three times in duplicate except starred (\*) were done twice in duplicate.

**Figure 4.**
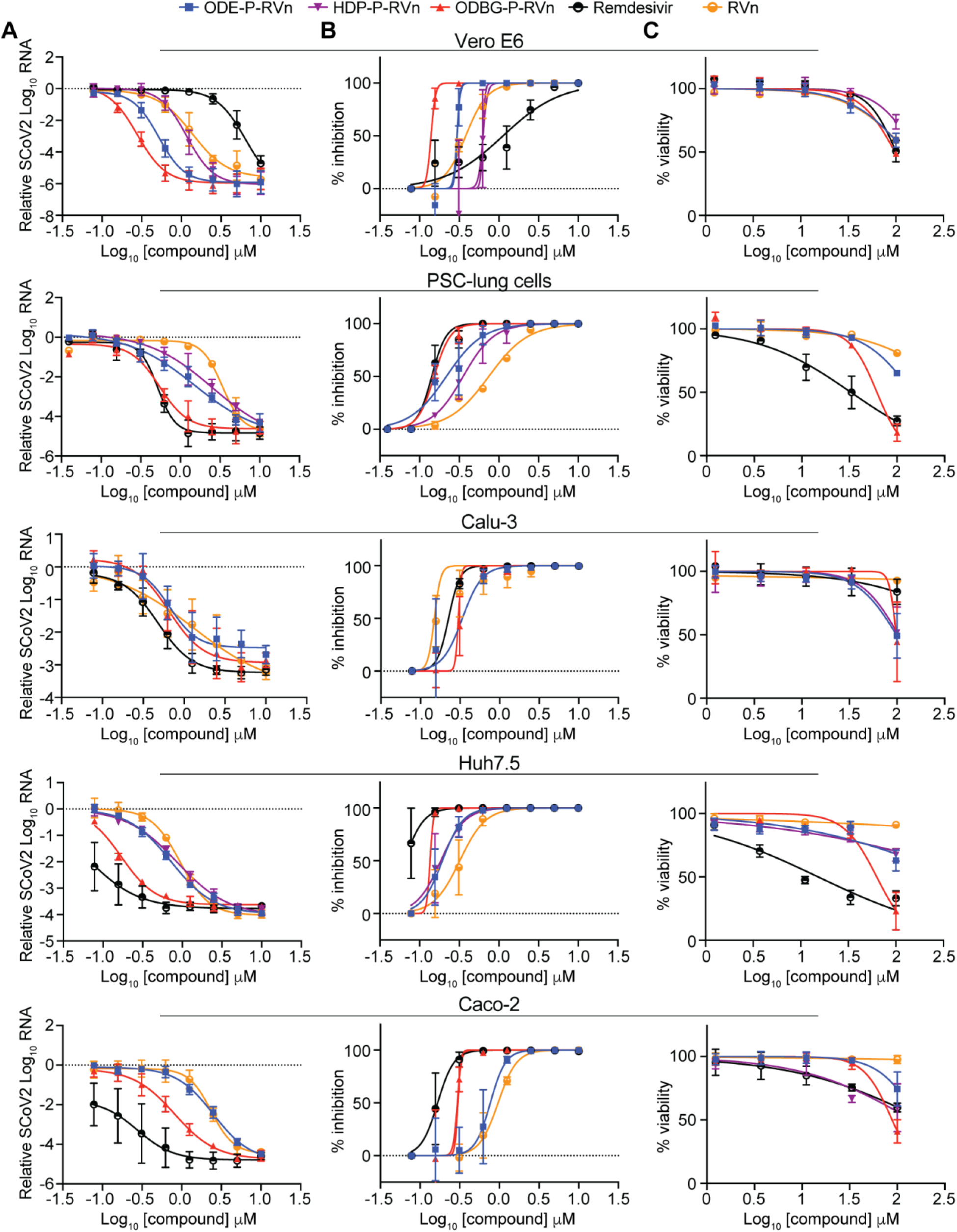
SARS-CoV-2 inhibitory activity replicate experiments. (A-B) Antiviral dose response curves for three Remdesivir analogs, Remdesivir (GS-5734), and Remdesivir nucleoside (GS-441524) against SARS-CoV-2 infection in multiple cell types. SARS-CoV-2 relative viral RNA reduction (A) and percent inhibition (B) in the indicated cell types. Cells were pretreated with the indicated dose of the indicated drug for 30 minutes and then infected with SARS-CoV-2 isolate USA-WA1/2020 for 48 hours. The relative SARS-CoV-2 Spike RNA expression was determined by qRT-PCR. (C) Cytotoxicity in the indicated cells incubated in the presence of the indicated drug at the indicated concentration for 48 hours, after which cell viability was measured by the CellTiter-Glo assay. Each (A-C) antiviral and cytotoxicity dose-response data point indicates the averages from 3 independent experiments performed in duplicate except as indicated in Table 1. Error bars represent the standard error mean (SEM).

We next measured the cytotoxicity of each compound by incubating each of these cell lines with serial dilutions of each compound from 1.23μM to 100μM for 48 hours (Figure 4C). The average 50% cytotoxic concentrations (CC_50_) for all compounds were greater than 60μM in all cell lines except for RDV which had a CC_50_ of 32.7μM in PSC-lung cells and 15.2μM in Huh7.5, a human hepatocyte cell line (Figure 4C and Table 1). The selectivity index of ODBG-P-RV ranged from 295 to 699 in the five cell types tested Table 1). The range of antiviral activity and cytotoxicity of ODBG-P-RVn (EC_50_ 0.14μM – 0.30μM and CC_50_ 61.5μM – 98.2μM) was more consistent across cell types than RDV (EC_50_ 0.06μM – 1.13μM and CC_50_ 15.2μM – >100μM) (Figure 5A – 5C and Table 1). Collectively, these data demonstrate that lipid RVn monophosphate prodrugs are potent antivirals against SARS-CoV-2 *in vitro* with low toxicity and excellent selectivity indexes.

**Figure 5.**
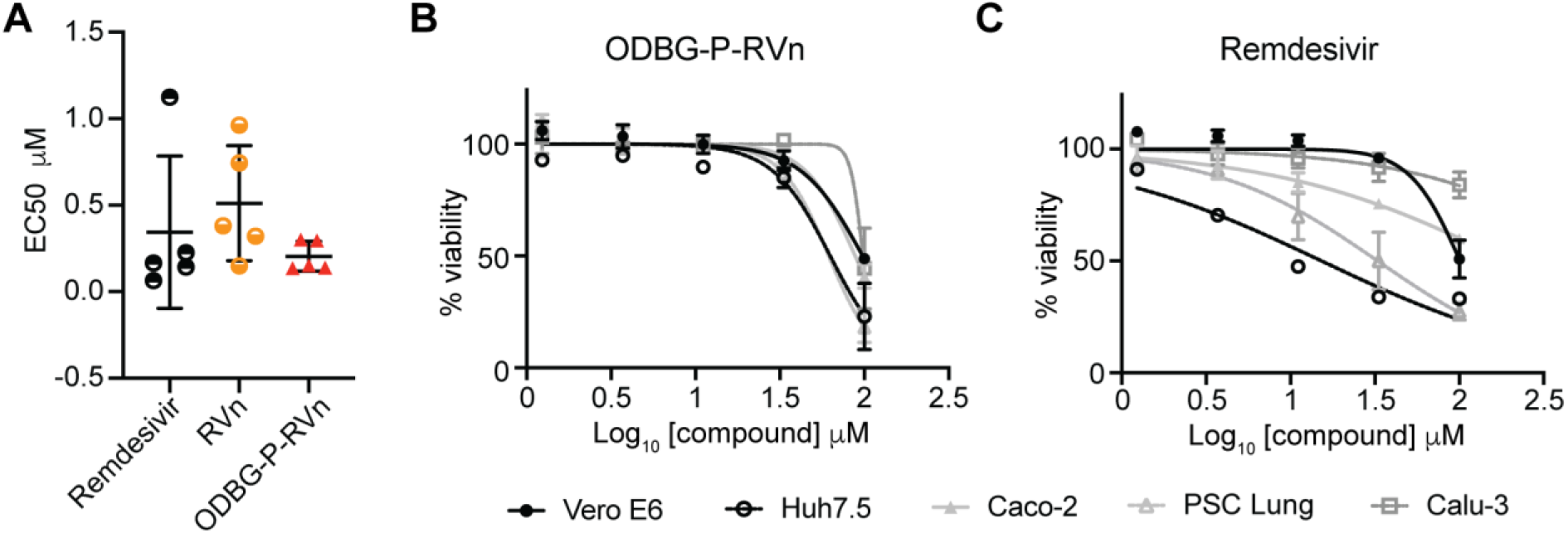
ODBG-P-RVn antiviral activity and cytotoxicity is highly reproducible across cell lines. (A) EC_50_ values for RDV, RVn and ODBG-P-RVn in 5 different cell lines derived from lung (human PSC-derived, Calu-3), kidney (Vero E6), colon (Caco-2), or liver (Huh7.5). (B-C) Cytotoxicity of (B) ODBG-P-RVn or (C) RDV in the indicated cells at the indicated concentration for 48 hours, as measured by the CellTiter-Glo assay.

### ODBG-P-RVn inhibits the human *Alphacoronavirus* 229E

To determine if ODBG-P-RVn inhibits other human coronaviruses, we performed antiviral assays in the human lung fibroblast cell line MRC-5 using a clinical isolate of human coronavirus 229E. MRC-5 cells were infected with 229E for 2 hours followed by incubation in the presence of varying concentrations of RDV, ODBG-P-RVn or vehicle for 72 hours post-infection. Both ODBG-P-RVn and RDV demonstrated a dose-dependent inhibition of cytopathic effect (CPE). The EC_50_ values of ODBG-P-RVn and RDV were 0.15μM and 0.04μM and the EC_90_s were 0.54 μM and 0.26 μM respectively (Figure 6A). The CC_50_ for ODBG-P-RVn and RDV were greater than 50μM in MRC-5 cells (Figure 6B). Together with the antiviral data for SARS-CoV-2, this demonstrates that ODBG-P-RVn has antiviral activity against two genetically distinct human pathogenic coronaviruses.

**Figure 6:**
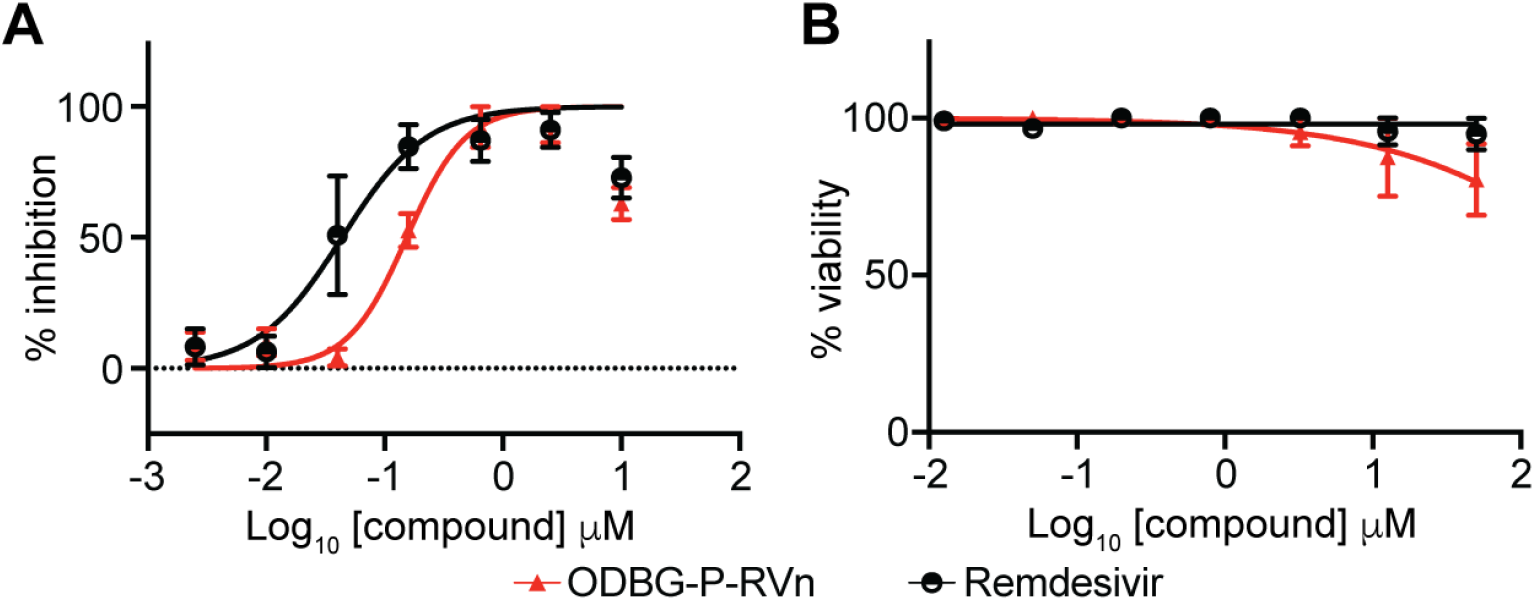
ODBG-P-RVn inhibits the human *Alphacoronavirus* 229E. (A) Antiviral dose response curves for Remdesivir (GS-5734) and ODBG-P-RVn against the human coronavirus 229E in MRC-5 cells. Cells were infected with 229E for 2 hours followed by treatment with the indicated dose of the indicated drug for 72 hours. The relative CPE was determined by measuring cell viability using an MTT assay. (B) Cytotoxicity in MRC-5 cells incubated in the presence of the indicated drug at the indicated concentration for 72 hours, after which cell viability was measured by the CellTiter-Glo assay. Data points indicate the averages from 3 independent experiments performed in duplicate. Error bars represent the standard error mean (SEM).

### Orally administered ODBG-P-RVn achieves therapeutic plasma levels in Syrian Hamsters

ODBG-P-RVn administered to Syrian Hamsters by oral gavage every 12 hours for seven days was well tolerated and no adverse clinical signs were noted. Peak plasma levels of ODBG-P-RVn were noted at 1 hour and fell by 50% in about 5 hours (Figure 7A). Plasma curves were generally similar at day 1 and 7 except at 16.9 mg/kg, the 7 day values were slightly higher than the levels at day 1. At 12 hours ODBG-P-RVn levels were above the EC_90_ for ODBG-P-RVn in all cell lines studied including Vero E6 cells and PSC lung cells on both day 1 and 7 (Figure 7A). Levels of the RVn, the nucleoside metabolite of ODBG-P-RVn peaked at 3 hours after administration and declined thereafter (Figure 7B). Plasma levels of RVn were less than the EC_90_ for RVn in both PSC lung cells and Vero E6 cells (Figure 7B). The observed low levels or RVn suggest that antiviral activity attributable to this metabolite will be minimal and are also consistent with finding of OBDG-P-RVn stability in human plasma (Figure 3). Collectively, these results suggest that ODBG-P-RVn will be effective in suppressing viral replication in a variety of tissue types *in vivo*.

**Figure 7.**
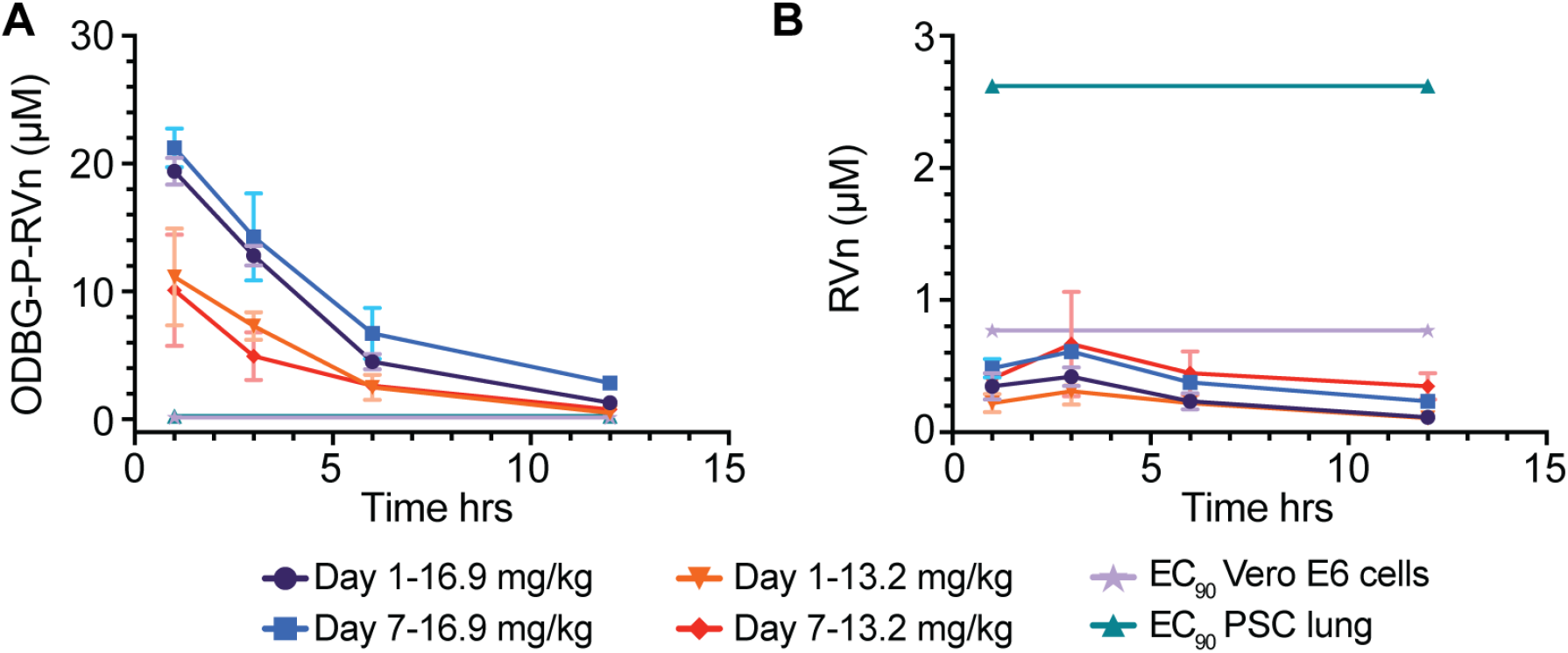
Seven Day Oral Pharmacokinetics in Syrian Hamsters. Syrian hamsters were given vehicle or ODBG-P-RVn by oral gavage every 12 hours for 7 days. Groups of 3 animals received vehicle or drug at doses of 16.9 and 13.2 mg/kg. Animals were weighed daily and monitored for clinical signs. Plasma samples were obtained at 1, 3, 6 and 12 hours on day 1 and day 7 and frozen for analysis of (A) ODBG-P-RVn and (B) RVn by LC/MS/MS.

## DISCUSSION

RDV is a prodrug designed to bypass the first phosphorylation of the Remdesivir nucleoside (RVn) which may be rate-limiting in the synthesis of RVn-triphosphate, the active metabolite. This occurs by the successive action of carboxyesterases, cathepsin A and phosphoramidases [16,25]. However, this approach does not appear to provide any benefit in Vero E6 cells, a monkey kidney cell line, as shown by Pruijssers et al [26] and by our results showing the antiviral activity of RVn is greater than that of RDV. Other perceived disadvantages of RDV include a lack of oral bioavailability, a difficult synthesis, instability in plasma, inadequate delivery to lung and hepatotoxicity. [13,16] In patients with Covid-19 and in the Syrian hamster model of SARS-CoV-2 disease, while high viral loads are notably present in the nasal turbinate, trachea and lung, as the infection proceeds, many other tissues also become infected, including the intestine, heart, liver, spleen, kidney, brain, lymph nodes and vascular endothelium. [27–31] However, RDV antiviral activity varies widely in lung and kidney cell lines with EC_50_ values of 1.65 μM in Vero E6 cells, 0.28 μM in Calu3 2B4, 0.010 μM in human alveolar epithelial cells (HAE), a 165-fold difference. [26] We found similar results in Vero E6 (EC_50_ 1.13 μM) and Calu-3 (0.23 μM). It has been suggested that this is due to variable amounts of the enzymes which convert RDV to RVn. [13,16] The antiviral activity of ODBG-P-RVn is consistently high in all five cell types we tested (Figure 5a).

We chose to design prodrugs of RVn which could provide oral bioavailability because an effective oral drug would allow for much earlier treatment of persons diagnosed with SARS-CoV-2 infection when active viral replication is believed to be the key driver of the subsequent course of the illness. We accomplished this by constructing liponucleotides of RVn resembling lysophospholipids that are normally highly absorbed intact in the GI tract. [32,33] Liponucleotides of this type are not metabolized in plasma and gain rapid entry to the cell often exhibiting greatly increased antiviral activity. [34–37]. In contrast to the activation of RDV which requires four transformations, intracellular kinase bypass with ODBG-P-RVn generates the RVn monophosphate directly when the lipid ester moiety is cleaved in a single reaction catalyzed by acid phospholipase C [38,39] or acid sphingomyelinase (sphingomyelin phosphodiesterase I) (K. Sandhoff and K. Hostetler, unpublished, 2013). ODBG-P-RVn, is likely to deliver relatively more drug to lung and less to liver as shown previously in lethal mousepox infection because of its apparent route to the circulation via intestinal lymph rather that portal vein. [23,24] Finally, the synthesis of these lipid prodrugs is much simpler than RDV and is readily scalable.

The limitations of RDV including a lack of oral bioavailability, a difficult synthesis, instability in plasma with rapid conversion to RVn, a less active metabolite, inadequate delivery to lung and dose-limiting hepatotoxicity provide opportunities to improve its clinical utility. As reported here, we synthesized three lipid prodrugs of RVn which were highly active in five cell types infected with SARS-CoV-2. The most active compound, ODBG-P-RVn, is 8 times more active than RDV in Vero E6 cells and equivalent to RDV in four other cell types. The cytotoxicity of ODBG-P-RVn is generally lower then RDV and more consistent across 5 cell lines and is not selectively higher in Huh7.5 cells, a hepatocyte cell line (Figure 5). Selectivity indexes are excellent and range from 295 to 699 in the five cell lines studied. ODBG-P-RVn achieved therapeutic levels in Syrian Hamsters by twice daily oral gavage and was well tolerated over a seven-day period of administration. Collectively, our data support the development of lipid prodrugs of RVn as potent oral antivirals that could be used orally early in the course of COVID-19 to prevent serious disease requiring hospitalization.

## MATERIALS AND METHODS

### Chemistry

All reagents were of commercial quality and used without further purification unless indicated otherwise. Chromatographic purification was done using the flash method with silica gel 60 (EMD Chemicals, Inc., 230–400 mesh). ^1^H, ^13^C and ^31^P nuclear magnetic resonance (NMR) spectra were recorded on either a Varian VX-500 or a Varian HG-400 spectrometer and are reported in units of ppm relative to internal tetramethylsilane at 0.00 ppm. Electrospray ionization mass spectra (ESI-MS) were recorded on a Finnigan LCQDECA mass spectrometer at the small molecule facility in the Department of Chemistry at University of California, San Diego. Purity of the target compounds was characterized by high performance liquid chromatography (HPLC) using a Beckman Coulter System Gold chromatography system. The analytical column was Phenomenex Synergi™ Polar-RP (4.6 × 150 mm) equipped with a SecurityGuard™ protection column. Mobile phase A was 95% water/5% methanol and mobile phase B was 95% methanol/5% water. At a flow rate of 0.8 mL/min, gradient elution was as follows: 10% B (0–3 min.); 10–95% B (3–20 min.); 95% B (20–25 min.); 95% to 10% B (25–34 min.). Compounds were detected by ultraviolet light (UV) absorption at 274 nm. Purity of compounds was also assessed by thin layer chromatography (TLC) using Analtech silica gel-GF (250 μm) plates and the solvent system: CHCl3/MeOH/conc NH4OH/H2O (70:30:3:3 v/v). TLC results were visualized with UV light, phospray (Supelco, Bellefonte, PA, USA) and charring at 400 °C.

### Compounds

Remdesivir (GS-5734) and Remdesivir nucleoside (GS-441524) were purchased from AA Blocks (San Diego, CA and Mason-Chem (Palo Alto, CA), respectively. 1-O-octadecyl-2-O-benzyl-sn-glycerol was obtained from Bachem, Torrance, CA).

### Synthesis of HDP-P-RVn: 5a. ((2*R*,3*S*,4*R*,5*R*)-5-(4-aminopyrrolo[2,1-*f*][1,2,4]triazin-7-yl)-5-cyano-3,4-dihydroxytetrahydrofuran-2-yl)methyl (3-(hexadecyloxy)propyl) hydrogen phosphate

N,N-Dicyclohexylcarbodiimide (DCC, 619 mg, 3 mmol) was added to a mixture of **2a** (300 mg, 0.91 mmol, prepared as in Warren et al [19], HDP-phosphate (**4a**, 414 mg, 1.10 mmol, prepared as in Kim et al [40], and 4-dimethylaminopyridine (DMAP, 122 mg, 1.0 mmol) in 25 mL of dry pyridine, and then the mixture was heated to 90 °C and stirred for 24h. Pyridine was then evaporated and the residue was purified by flash column chromatography on silica gel 60. Gradient elution (CH_2_Cl_2_/methanol 10-20%) afforded 423 mg (67% yield) of 2’,3’-isopropylidene derivative of **5a**. ^1^H NMR (500 MHz, chloroform-*d*) δ 8.42 (s, 1H), 7.98 (s, 1H), 7.70 (s, 2H), 6.22 (d, *J* = 6.0 Hz, 1H), 5.68 (d, *J* = 6.2 Hz, 1H), 5.15 (d, *J* = 1.0 Hz, 1H), 4.70 (dd, *J* = 3.8, 0.9 Hz, 1H), 4.48 – 4.42 (m, 1H), 4.26 (ddd, *J* = 11.2, 8.5, 2.6 Hz, 1H), 4.15 (ddd, *J* = 11.1, 8.5, 2.6 Hz, 1H), 4.02 (dt, *J* = 8.5, 6.3 Hz, 2H), 3.49 (t, *J* = 6.1 Hz, 2H), 3.40 (t, *J* = 6.1 Hz, 2H), 1.95 (p, *J* = 6.2 Hz, 2H), 1.54 (tt, *J* = 7.4, 6.1 Hz, 2H), 1.31 (s, 3H), 1.32 – 1.24 (m, 26H), 0.94 – 0.85 (m, 3H). ESI MS 691.6 [M-H]^-^.

Concentrated HCl (0.1 mL) in tetrahydrofuran (THF) was added to a stirred solution of 2’,3’-isopropylidene-**5a** (100 mg, 0.14 mmol) in THF (10 mL) at room temperature. The mixture was stirred for 3h and then sodium bicarbonate (50 mg) and water (2 mL) were added. After stirring an additional 15 min. the solvents were evaporated and cold water (10 mL) was added to the residue. The solid product was collected by vacuum filtration and dried under vacuum to yield compound **5a** (79 mg, 87% yield) as an off-white solid. ^1^H NMR (500 MHz, CDCl_3_-methanol-*d*_4_) *δ* ppm ^1^H NMR (500 MHz, Chloroform-*d*) δ 8.42 (s, 1H), 7.98 (s, 1H), 7.70 (s, 1H), 6.22 (d, *J* = 6.0 Hz, 1H), 5.70 (d, *J* = 6.0 Hz, 1H), 5.12 (d, *J* = 4.2 Hz, 1H), 4.55 (ddd, *J* = 5.5, 2.7, 0.9 Hz, 1H), 4.40 (dtd, *J* = 6.8, 2.6, 0.8 Hz, 1H), 4.33 – 4.27 (m, 2H), 4.25 (ddd, *J* = 11.1, 8.4, 2.6 Hz, 1H), 4.16 (ddd, *J* = 11.3, 8.5, 2.6 Hz, 1H), 4.02 (dt, *J* = 8.5, 6.3 Hz, 2H), 3.49 (t, *J* = 6.1 Hz, 2H), 3.40 (t, *J* = 6.1 Hz, 2H), 1.95 (p, *J* = 6.2 Hz, 2H), 1.59 – 1.50 (m, 1H), 1.34 – 1.24 (m, 23H), 0.94 – 0.85 (m, 3H). ESI MS: 652.39 [M-H]^-^. Purity by HPLC: 99.7%

### Synthesis of ODE-P-RVn, 5b. ((2*R*,3*S*,4*R*,5*R*)-5-(4-aminopyrrolo[2,1-*f*][1,2,4]triazin-7-yl)-5-cyano-3,4-dihydroxytetrahydrofuran-2-yl)methyl (2-(octadecyloxy)ethyl) hydrogen phosphate

N,N-Dicyclohexylcarbodiimide (DCC, 0.3 g, 1.4 mmol) was added to a mixture of **2a** (0.23 g, 0.7 mmol), ODE-phosphate (**4b**, 0.27 g, 0.68 mmol), and 4-dimethylaminopyridine (DMAP, 0.07 g, 0.6 mmol) in 10 mL of dry pyridine, and then the mixture was heated to 90 °C and stirred for 3 days. Pyridine was then evaporated and the residue was purified by flash column chromatography on silica gel 60. Gradient elution (CH_2_Cl_2_/methanol 10-20%) afforded 0.22 g (45% yield) of 2’,3’-isopropylidene-**5b**. Concentrated HCl (0.3 mL) was added slowly to a stirred solution of 2’,3’-isopropylidene-**5b** (0.2 g, 0.28 mmol) in tetrahydrofuran (2 mL) at 0 °C. The mixture was allowed to warm to room temperature overnight and then was diluted with water (2 mL) and adjusted to pH = 8 by adding saturated sodium bicarbonate. The product was extracted with chloroform (3 x 30 mL) and the organic layer was concentrated under reduced pressure. The residue was purified by flash chromatography on silica gel. Elution with 20% MeOH/CH_2_Cl_2_ gave 0.10 g (55% yield) of compound **5b**. ^1^H NMR (400 MHz, CDCl_3_-methanol-*d*_4_) *δ* ppm 7.89 (s, 1 H), 6.94 (d, *J*=4.65 Hz, 1H), 6.89 (d, *J*=4.65 Hz, 1H), 4.40 (d, *J*=4.65 Hz, 2H), 4.21 - 4.28 (m, 1H), 4.12 - 4.20 (m, 1H), 4.04 - 4.12 (m, 1H), 3.91 (d, *J*=4.89 Hz, 2H), 3.46 - 3.57 (m, 2H), 3.42 (td, *J*=6.85, 1.96 Hz, 2H), 3.34 (dt, *J*=3.18, 1.59 Hz, 2H), 1.53 (d, *J*=6.85 Hz, 2H), 1.20 - 1.37 (m, 30H), 0.89 (t, *J*=6.97 Hz, 3H). ESI MS: 666.43 [M-H]^-^. Purity by HPLC 98.4%.

### Synthesis of ODBG-P-RVn, 5c. ((2*R*,3*S*,4*R*,5*R*)-5-(4-aminopyrrolo[2,1-*f*][1,2,4]triazin-7-yl)-5-cyano-3,4-dihydroxytetrahydrofuran-2-yl)methyl ((*R*)-2-(benzyloxy)-3-(octadecyloxy)propyl) hydrogen phosphate

N,N-Dicyclohexylcarbodiimide (DCC, 310 mg, 1.5 mmol) was added to a mixture of **2a** (300 mg, 0.91 mmol), ODBG-phosphate (**4c**, 515 mg, 1.0 mmol), and 4-dimethylaminopyridine (DMAP, 122 mg, 1.0 mmol) in 25 mL of dry pyridine, and then the mixture was heated to 90 °C and stirred for 24h. Pyridine was then evaporated and the residue was purified by flash column chromatography on silica gel 60. Gradient elution (CH_2_Cl_2_/methanol 10-20%) afforded 210 mg (28% yield) of compound 2’,3’-isopropylidene-**5c**. ESI MS 826.58 [M-H]^-^. Concentrated HCl (0.1 mL) in tetrahydrofuran (THF) was added to a stirred solution of 2’,3’-isopropylidene-**5c** (210 mg, 0.25 mmol) in THF(10 mL) at room temperature. The mixture was stirred for 3h and then sodium bicarbonate (50 mg) and water (2 mL) were added. After stirring an additional 15 min. the solvents were evaporated and cold water (10 mL) was added to the residue. The solid product was collected by vacuum filtration and dried under vacuum to yield compound **5c** (71 mg, 36% yield) as an off-white solid. ^1^H NMR (500 MHz, DMSO-d_6_) δ 7.89 (s, 1H), 7.79 (s, 1H), 7.33 – 7.24 (m, 4H), 7.22 (ddd, J = 8.7, 5.3, 2.5 Hz, 1H), 6.88 (d, J = 4.5 Hz, 1H), 6.80 (d, J = 4.6 Hz, 1H), 6.14 (d, J = 5.2 Hz, 1H), 5.91 (s, 1H), 4.55 (q, J = 12.1, 12.1, 12.1 Hz, 3H), 4.10 (dt, J = 6.7, 4.3, 4.3 Hz, 1H), 3.93 (t, J = 5.9, 5.9 Hz, 1H), 3.79(dddd, J = 28.2, 12.1, 7.9, 4.4 Hz, 2H), 3.62 (tdd, J = 10.9, 10.9, 8.2, 5.1 Hz, 4H), 3.43 (dd, J = 10.6, 3.5 Hz, 2H), 1.43 (p, J = 6.6, 6.6, 6.6, 6.6 Hz, 2H), 1.21 (d, J = 8.3 Hz, 30H), 0.83 (t, J = 7.0, 7.0 Hz, 3H). ESI MS: 786.48 [M-H]^-^. Purity by HPLC: 97.6%.

### Cells

Vero E6, Caco-2, and Calu-3 cell lines were obtained from ATCC. Huh7.5 cells were obtained from Apath LLC. Calu-3 and Caco-2 cells were propagated in MEM (Corning), 10% FBS, Penicillin-Streptomycin (Gibco). Vero E6 and Huh7.5 cells were propagated in DMEM (Corning) with 10% FBS and Penicillin-Streptomycin (Gibco). For human PSC-lung cell generation, human lung organoids were generated as previously described. [41] H9 embryonic stem cells (WiCell) were cultured in feeder free conditions upon Matrigel (Corning #354230) coated plates in mTeSR medium (StemCellTech #85850). Media was changed daily, and stem cells were passaged using enzyme free dissociation reagent ReLeSR™ (Stem Cell Tech#05872). Cultures were maintained in an undifferentiated state, in a 5% CO2 incubator at 37 °C. For proximal lung organoid generation, human PSCs were dissociated into single cells, and then seeded on Matrigel-coated plates (BD Biosciences) at a density of 5.3 x 10^4^ cells/cm^2^ in Definitive Endoderm (DE) induction medium (RPMI1640, 2% B27 supplement, 1% HEPES, 1% glutamax, 50 U/mL penicillin/streptomycin), supplemented with 100 ng/mL human activin A (R&D), 5μM CHIR99021 (Stemgent), and 10μM ROCK inhibitor, Y-27632 (R&D Systems) on day 1. On days 2 and 3 cells were cultured in DE induction media with only 100 ng/mL human activin A. Anterior Foregut Endoderm (AFE) was generated by supplementing serum free basal medium (3 parts IMDM:1 part F12, B27+N2 supplements, 50U/mL penicillin/streptomycin, 0.25% BSA, 0.05 mg/mL L-ascorbic acid, 0.4mM monothioglycerol) with 10μM SB431542 (R&D) and 2 μM Dorsomorphin (StemGent) on days 4-6. On day 7, AFE medium was changed to Lung Progenitor Cell (LPC) induction medium, containing serum free basal medium supplemented with 10 ng/mL human recombinant BMP4 (R&D), 0.1 μM all-trans retinoic acid (Sigma-Aldrich) and 3μM CHIR99021. Media was changed every other day for 9-11 days. To generate 3D human proximal lung organoids, we modified a previously published protocol. [42] LPCs were dissociated in accutase for 10minutes and resuspended in Matrigel in a 12-well, 0.4μm pore size Transwell (Corning) culture insert at 5.0 x 10^4^ cells/200ul of Matrigel. Cells were cultured in proximal lung organoid maturation media using serum free basal medium supplemented with 250ng/mL FGF2, 100ng/mL rhFGF10, 50nM dexamethasone (Dex), 100μM 8-Bromoadenosine 3’,5’-cyclic monophosphate sodium salt (Br-cAMP), 100μM 3-Isobutyl-1-methylxanthine (IBMX) and 10 μM ROCK inhibitor (Y-27632). Proximal lung organoid media was changed every other day for 3 weeks. Human PSC-derived lung organoids were dissociated into single cells and seeded at 20,000 cells per well of a matrigel coated 96-well plate one day before transfection. Transwells containing the proximal organoids in matrigel were incubated in 2U/ml dispase for 30 minutes at 37 °C. Cold PBS was added to the mixture then centrifuged at 400 x g for 5 mins. Supernatant was carefully removed and resuspended in 2-3mls of TrypLE Express (Gibco # 12605010) for 20 minutes at 37 °C. Reaction was quenched with 2% FBS in DMEM/F12 then centrifuged at 400 x g for 5 min. The supernatant was aspirated, and the cell pellet resuspended in 1ml of quenching media supplemented with 10 μM Rock inhibitor (Y-27632). Cell count was performed and the respective volume of cells were transferred into a reagent reservoir trough and resuspended in proximal lung organoid maturation media and plated via multichannel pipette into 96 well plates at 100ul per well as monolayers.

### SARS-CoV-2 infection

SARS-CoV-2 isolate USA-WA1/2020 (BEI Resources) was propagated and infectious units quantified by plaque assay using Vero E6 (ATCC) cells. Approximately 12,000 cells from each cell line were seeded per well in a 96 well plate. Vero E6 and Huh7.5 were seeded approximately 24h prior to treatment/infection. Calu-3 and Caco-2 were seeded approximately 48h prior to treatment/infection. Human PSC lung cell infections and cytotoxicity experiments were performed when cells reached 100% confluency. Compounds or controls were added at the indicated concentrations 30 minutes prior to infection followed by the addition of SARS-CoV-2 at a multiplicity of infection equal to 0.01. After incubation for 48 hours at 37°C and 5% CO_2_, cells were washed twice with PBS and lysed in 200ul TRIzol (ThermoFisher). All work with SARS-CoV-2 was conducted in Biosafety Level 3 conditions at the University of California San Diego with approval from the Institutional Biosafety Committee.

### Human Coronavirus 229E infection

Human Coronavirus 229E (ATCC) was propagated and infectious units quantified by TCID_50_ using MRC-5 cells. For antiviral testing, approximately 10^4^ MRC-5 cells were seeded per well in EMEM (10%FCS) at 37C in a 96 well plate overnight. Medium from each well was removed and cells were infected with 100 TCID_50_ virus in 100 μL medium for two hours. Cells were washed one time with medium and then compounds or controls added at the indicated concentrations. After three days, CPE was observed under microscope and quantified using an MTT cell proliferation assay kit (Abcam) read on an ELx800, Universal Microplate reader (BIO-TEK Instruments, INC). The % Inhibition was calculated as (A_tv_ – A_cv_)/(A_cd_ – A_cv_) × 100% where A_tv_ indicates the absorbance of the test compounds with virus infected cells and A_cv_ and A_cd_ indicate the absorbance of the virus control and the absorbance of the cell control, respectively. The average half-maximal effective concentration (EC_50_) was defined as the concentration which achieved 50% inhibition of virus-induced cytopathic effects.

### RNA extraction, cDNA synthesis and qPCR

RNA was purified from TRIzol lysates using Direct-zol RNA Microprep kits (Zymo Research) according to manufacturer recommendations that included DNase treatment. RNA was converted to cDNA using the iScript cDNA synthesis kit (BioRad) and qPCR was performed using iTaq universal SYBR green supermix (BioRad) and an ABI 7300 real-time pcr system. cDNA was amplified using the following primers RPLP0 F – GTGTTCGACAATGGCAGCAT; RPLP0 R – GACACCCTCCAGGAAGCGA; SARS-CoV-2 Spike F – CCTACTAAATTAAATGATCTCTGCTTTACT; SARS-CoV-2 Spike R – CAAGCTATAACGCAGCCTGTA. Relative expression of SARS-CoV-2 Spike RNA was calculated by delta-delta-Ct by first normalizing to the housekeeping gene RPLP0 and then comparing to SARS-CoV-2 infected Vero E6 cells that were untreated (reference control). Curves were fit using the nonlinear regression – log(inhibitor) vs. response (four parameter) model using Prism 9. To calculate effective concentrations EC_50_ and EC_90_ values, qRT-PCR values were normalized to percent inhibition and curves fit using the nonlinear regression – log(agonist) vs. response (four parameter) model with bottom and top constrained to 0 and 100 respectively using Prism 9.

### Cell viability assay

Cell type were seeded as per SARS-CoV-2 infection studies in opaque walled 96-well cell culture plates or 229E infection studies in clear 96-well cell culture plates and incubated overnight. Compounds or controls were added at the indicated concentrations. For SARS-CoV-2 related studies, cells were incubated for 48.5 hours at 37°C and 5% CO_2_, an equal volume of CellTiter-Glo reagent (Cat. # G7570, Promega, Madison, WI) was added, mixed and luminescence recorded on a Veritas Microplate Luminometer (Turner BioSystems) according to manufacturer recommendations. For 229E related, cells were incubated for 72 hours at 37°C and 5% CO_2_, supernatants removed, 50μL of serum-free media and 50μL of MTT Reagent (Abcam ab211091) added to each well and incubated for 3hrs at 37C. Absorbance was measured on an ELx800, Universal Microplate reader, (BIO-TEK Instruments, INC) according to manufacturer recommendations. Percent viability was calculated compared to untreated controls and CC_50_ values were calculated using Prism 9.

### Stability in human plasma

ODBG-P-RVn was incubated in human plasma with K_2_EDTA or sodium heparin as the anticoagulant. The final concentration of ODBG-P-RVn was 2.00 μg/mL in the incubation. After incubation at 37°C, samples of the test article incubations were taken at 0 (pre-incubation), 0.5, 1, 2, 4, 8 and 24 hours and immediately frozen at −70°C. The extracts were prepared by a solid phase extraction using a Waters Sep Pak® tC18 25 mg SPE plate and analyzed as below.

### Analytical Methods: ODBG-P-RVn

Hamster plasma samples (10 μL) containing ODBG-P-RVn and K_2_EDTA as the anticoagulant were added to polypropylene tubes containing water (100 μL), internal standard solution (10 μL; 1,000 ng/mL of ODE-P-RVn in ACN:DMF (1:1, v/v)), and 10 μL of ACN:DMF (1:1, v/v). The solutions were mixed, then acidified with phosphoric acid, 85% w/v:water (1:19, v/v; 10 μL), mixed, then diluted with 200 μL of IPA, mixed, then diluted with 500 μL of water, and mixed. The samples were extracted with a Sep-Pak® tC18 96-well solid phase extraction plate (25 mg; Waters, Milford, MA). Extraction occurred under positive pressure conditions using nitrogen. Samples were washed serially with 1 mL of water:acetonitrile:formic acid (475:25:0.5, v/v/v) and 0.4 mL of water:acetonitrile:formic acid (350:150:0.5, v/v/v) before being serially eluted with 100 μL and 150 μL of water:{acetonitrile:isopropyl alcohol (1:1, v/v)}:formic acid:ammonium formate:citric acid solution, 2% w/v (15:85:0.1:0.1:0.1,v/v/v/w/v). The citric acid solution was prepared as water:citric acid monohydrate (20:0.4, v/w). After elution, 100 μL of water was added to each sample. The ODBG-P-RVn extracts were analyzed using an Agilent 1200 HPLC system (Agilent, Santa Clara, CA) coupled to an API5500 mass analyzer (SCIEX, Foster City, CA). Analytes were chromatographically separated using a Dacapo DX-C18 MF column (100 x 2 mm, 2.5 μm; ImtaktUSA, Portland, OR) using a mobile phase system consisting of Mobile Phase A (water: formic acid:[water:ammonium formate:citric acid (25:5:0.5, v/w/w)] (1,000:1:1, v/v/v) and Mobile Phase B (acetonitrile:isopropyl alcohol:formic acid:[water:ammonium formate:citric acid (25:5:0.5, v/w/w)] (800:200:1:1, v/v/v/v). The total analytical run time was 4.5 minutes. The mobile phase was nebulized using heated nitrogen in a Turbo-V source/interface set to electrospray positive ionization mode. The ionized compounds were detected using multiple reaction monitoring with transitions m/z 788.4 > 229 (V2043) and 668.4 > 467.2 (V2041). This method is applicable for measuring ODBG-P-RVn concentrations ranging from 6.25 to 3,000 ng/mL using 10.0 μL of plasma for extraction. The peak areas of ODBG-P-RVn and RVn were acquired using Analyst v. 1.6.2 (SCIEX, Framingham, MA). The calibration curve was obtained by fitting the peak area ratios of the analyte/I.S. and the standard concentrations to a linear equation with 1/x2 weighting, using Analyst. The equation of the calibration curve was then used to interpolate the concentrations of the analyte in the samples using their peak area ratios. The peak areas used for the calculations were not rounded.

### RVn (GS441524)

Hamster plasma samples (20 *μ*L) containing GS-441524 and K_2_EDTA as the anticoagulant were added to Eppendorf LoBind microfuge tubes containing acetonitrile (300 *μ*L) and water:acetonitrile (2:8, v/v; 60 *μ*L). The solutions were mixed and centrifuged at 16,000 g for five minutes. The supernatant (300 *μ*L) was then filtered through an Ostro protein precipitation and phospholipid removal plate (25 mg; Waters, Milford, MA). Filtration occurred under positive pressure conditions using nitrogen. Collected filtered samples were capped, mixed and stored at 10°C pending analysis. The GS-441524 extracts were analyzed using an Acquity UPLC system (Waters, Milford, MA) coupled to a G2-S QTof mass analyzer (Waters, Milford, MA). Analytes were chromatographically separated using a Unison-UK Amino HT column (100 x 2 mm, 3 μm; ImtaktUSA, Portland, OR) using a mobile phase system consisting of Mobile Phase A (0.008% ammonium hydroxide, 0.012% acetic acid in water, v/v/v) and Mobile Phase B (0.008% ammonium hydroxide, 0.012% acetic acid in acetonitrile, v/v/v). The total analytical run time was 12.5 minutes. The mobile phase was nebulized using heated nitrogen in a Z-spray source/interface set to electrospray positive ionization mode. The ionized compounds were detected using Tof MS scan monitoring in sensitivity mode scanning from 50.0 to 700 *m/z*. This method is applicable for measuring GS-441524 concentrations ranging from 1.00 to 1,000 ng/mL using *μ*L of plasma for extraction. The peak areas of GS-441524 were acquired using MassLynx V4.2 (Waters, Milford, MA). The calibration curve was obtained by fitting the peak area ratios of the analyte and the standard concentrations to a linear equation with 1/x^2^ weighting using MassLynx. The equation of the calibration curve was then used to interpolate the concentrations of the analyte in the samples using their peak areas. The peak areas used for the calculations were not rounded.

## ACKNOWLEDGEMENTS

This research was supported by NIAID grant RO1 AI131424, the San Diego Center for AIDS Research, NIH grant (K08 AI130381) and Career Award for Medical Scientists from the Burroughs Welcome Fund to AFC, and by CIRM (DISC2COVID19-12022) to SLL. The following reagent was deposited by the Centers for Disease Control and Prevention and obtained through BEI Resources, NIAID, NIH: SARS-Related Coronavirus 2, Isolate USA-WA1/2020, NR-52281.

